# Frequency-independent biological signal identification (FIBSI): A free program that simplifies intensive analysis of non-stationary time series data

**DOI:** 10.1101/2020.05.29.123042

**Authors:** Ryan M. Cassidy, Alexis G. Bavencoffe, Elia R. Lopez, Sai S. Cheruvu, Edgar T. Walters, Rosa A. Uribe, Anne Marie Krachler, Max A. Odem

## Abstract

Extracting biological signals from non-linear, dynamic and stochastic experimental data can be challenging, especially when the signal is non-stationary. Many currently available methods make assumptions about the data structure (e.g., signal is periodic, sufficient recording time) and modify the raw data in pre-processing using filters and/or transformations. With an agnostic approach to biological data analysis as a goal, we implemented a signal detection algorithm in Python that quantifies the dimensional properties of waveform deviations from baseline via a running fit function. We call the resulting free program frequency-independent biological signal identification (FIBSI). We demonstrate the utility of FIBSI on two disparate types of experimental data: *in vitro* whole-cell current-clamp electrophysiological recordings of rodent sensory neurons (i.e., nociceptors) and *in vivo* fluorescence image time-lapse movies capturing gastrointestinal motility in larval zebrafish. In rodent nociceptors, depolarizing fluctuations in membrane potential are irregular in shape and difficult to distinguish from noise. Using FIBSI, we determined that nociceptors from naïve mice generate larger, more frequent fluctuations compared to naïve rats, suggesting species-specific specializations in rodent nociceptors. In zebrafish, measuring gut motility is a useful tool for addressing developmental and disease-related mechanisms associated with gut function. However, available methods are laborious, technically complex, and/or not cost-effective. We developed and tested a novel assay that can characterize intestinal peristalsis using imaging time series datasets. We used FIBSI to identify muscle contractions in the fluorescence signals and compared their frequencies in unfed and fed larvae. Additionally, FIBSI allowed us to discriminate between peristalsis and oscillatory sphincter-like movements in functionally distinct gut segments (foregut, midgut, and cloaca). We conclude that FIBSI, which is freely available via GitHub, is widely useful for the unbiased analysis of non-stationary signals and extraction of biologically meaningful information from experimental time series data and can be employed for both descriptive and hypothesis-driven investigations.

**Author Summary:** Biologists increasingly work with large, complex experimental datasets. Those datasets often encode biologically meaningful signals along with background noise that is recorded along with the biological data during experiments. Background noise masks the real signal but originates from other sources, for example from the equipment used to perform the measurements or environmental disturbances. When it comes to analyzing the data, distinguishing between the real biological signals and the background noise can be very challenging. Many existing programs designed to help scientists with this problem are either difficult to use, not freely available, or only appropriate to use on very specific types of datasets. The research presented here embodies our goal of helping others to analyze their data by employing a powerful but novice-friendly program that describes multiple features of biological activity in its raw form without abstract transformations. We show the program’s applicability using two different kinds of biological activity measured in our labs. It is our hope that this will help others to analyze complex datasets more easily, thoroughly, and rigorously.

## 1. Introduction

It can be difficult to identify and characterize biological signals encoded within non-linear, dynamic and stochastic experimental data using analytical methods without introducing bias due to *a priori* assumptions about the data structure and nature of the signals. This is a challenge faced widely across biological disciplines as many biological datasets are non-ideal (e.g., the frequency resolution is low due to undersampling of the time domain, or the signal-to-noise ratio is low due to limited measurement sensitivity) and the encoded signals are often of irregular shape. The Fourier transform and derivative analytical tools [1], which are widely used to analyze biological time series, have limited capacity to accurately identify and characterize biological signals encoded in such data. A wealth of non-linear methods exist that aim at analyzing these kinds of data series (e.g., using filters and transformations [2,3], multifractals for invariant structures [4]), but most are field-specific and only applicable to a specific type of experimental data (e.g., electroencephalogram and electrocardiogram recordings [5,6]), and would require major optimization prior to application to other types of experimental data with different data structures and signal shapes. Information about a signal of interest is obtained from data that has been subject to differing degrees of pre- and post-hoc processing. We contend that biologically meaningful, easily interpretable information can be readily derived from signals in their raw form, even if they are non-stationary.

In this paper, we demonstrate the utility of a novel frequency-independent biological signal identification (FIBSI) program as a first-line tool for isolating biological signals from unprocessed (raw) time series data. We demonstrate the utility of FIBSI using two different types of experimental data series that were acquired using unrelated techniques: *in vitro* whole-cell current-clamp electrophysiological recordings of depolarizing spontaneous fluctuations (DSFs) of membrane potential from dissociated rodent putative C-type sensory neurons (primarily nociceptors) and *in vivo* fluorescence time-lapse imaging data of motility in the larval zebrafish intestine. Each type of data presents commonly encountered challenges that deter many researchers. The DSFs in nociceptors are irregular in waveform and occurrence, small in magnitude, and thus can be difficult to distinguish as a biologically relevant signal from internal and external noise in neuronal recordings (i.e., low signal-to-noise ratios). For the analysis of gut motility, the signal-to-noise ratios can vary substantially between regions of the intestinal tract and between experimental animals. Motility frequencies can also differ within and between regions within specimens. Challenges in quantifying fluorescence signals are similar to those encountered with identifying electrophysiological signals, with the additional hurdle of reliably identifying a region of interest (ROI) that shows the signal of interest without high dilution due to background noise. Previously published methods circumvent these issues by different means. For example, some focus on low frequency, high signal-to-noise ratio data collected from ROIs with substantial physical separation from each other [7,8], use post-hoc analyses with a second identification method (e.g., antibody staining or tight localization to neuronal nuclei) to tie ROIs to relevant physical locations [9,10], use a short stimulus and test window to isolate activity [11], and/or use phasic stimulations to identify ROIs and the Fourier transform to reduce noise [12–14]. In many systems, however, the choice of ROIs is not flexible, and ROIs containing the relevant biological signal of interest are intrinsically noisy, so the above-mentioned approaches are not applicable. Here we demonstrate how FIBSI overcomes these common challenges without modifications to the raw data and minimal ad-hoc assumptions. The signal (event) detection algorithm core to FIBSI adapts the Ramer-Douglas-Peucker algorithm [15,16] to break down waveforms above and below a time-dependent running fit function. The dimensional properties of the isolated signals (events) can then be used for descriptive and hypothesis-driven statistical analyses.

## 2. Results

### 2.1. FIBSI allows an unbiased comparative analysis of membrane potential fluctuations in rodent nociceptors

We have characterized two predominant types of small-diameter C-type sensory neurons in rodents using whole-cell current-clamp electrophysiology: rapidly-accommodating (RA) and non-accommodating (NA) neurons [17,18]. Here, we used a subset of RA and NA neurons dissociated from 16 male Sprague-Dawley rats and 1 male and 4 female C57BL/6 wild-type mice for post-hoc analysis. These rodents were used as naïve controls in prior studies performed by our research group [18,19]. No major differences in electrophysiological properties were reported between genders [18], so the neurons isolated from male and female mice were pooled. Like in previous studies, these neurons were distinguished by injecting 2 s pulses of current at increasing increments of 5 pA until each neuron reached rheobase by eliciting an action potential (AP). All RA neurons elicited a single AP at the beginning of the pulse (**Fig 1A**), while a majority of NA neurons exhibited delayed AP firing at rheobase (**Fig 1B-C**). At twice rheobase, RA neurons only fired a single AP at the beginning of the pulse (**Fig 1D**), while NA neurons elicited repetitive firing (**Fig 1E-F**). Additionally, both RA and NA neurons generated irregular, low-frequency DSFs in membrane potential. In NA neurons, the DSFs were notably larger in amplitude and those that reached threshold potential generated an AP. Prolonged depolarization to −45 mV increases the likelihood of rodent nociceptors to be active and generate DSFs, and is used as an experimental setting to model inflammatory and pain-like conditions *in vitro* [17,18]. However, no direct comparisons of the DSFs across rodent species have been performed. To make the experimental conditions comparable between neuron types and rodent species, we chose a subset of neurons that were depolarized to −45 mV for 30 s [17,18] for FIBSI analysis. Both RA and NA neurons are abundant in rats (~30% and ~70% respectively) [17], but RA neurons are sparse in mice (~6%) [18], and too few had been sampled to warrant further analysis. Rat RA neurons did not exhibit ongoing activity when depolarized to −45 mV, and DSFs were small in magnitude (**Fig 1G**). A fraction (~20%) of the rat and mouse NA neurons exhibited ongoing activity and larger DSFs (**Fig 1H-I**), as reported previously [17,18]. We also tested for other signs of increased excitability in NA neurons using standard membrane property measures. The mouse NA neurons had significantly lower membrane capacitances, lower rheobase values, and depolarized resting membrane potentials in comparison to rat NA neurons (**Fig 1J-L**), but their AP thresholds were similar (**Fig 1M**). In comparison to rat RA neurons, rat NA neurons had lower rheobase values and hyperpolarized AP thresholds, like reported previously [17].

**Fig 1.**
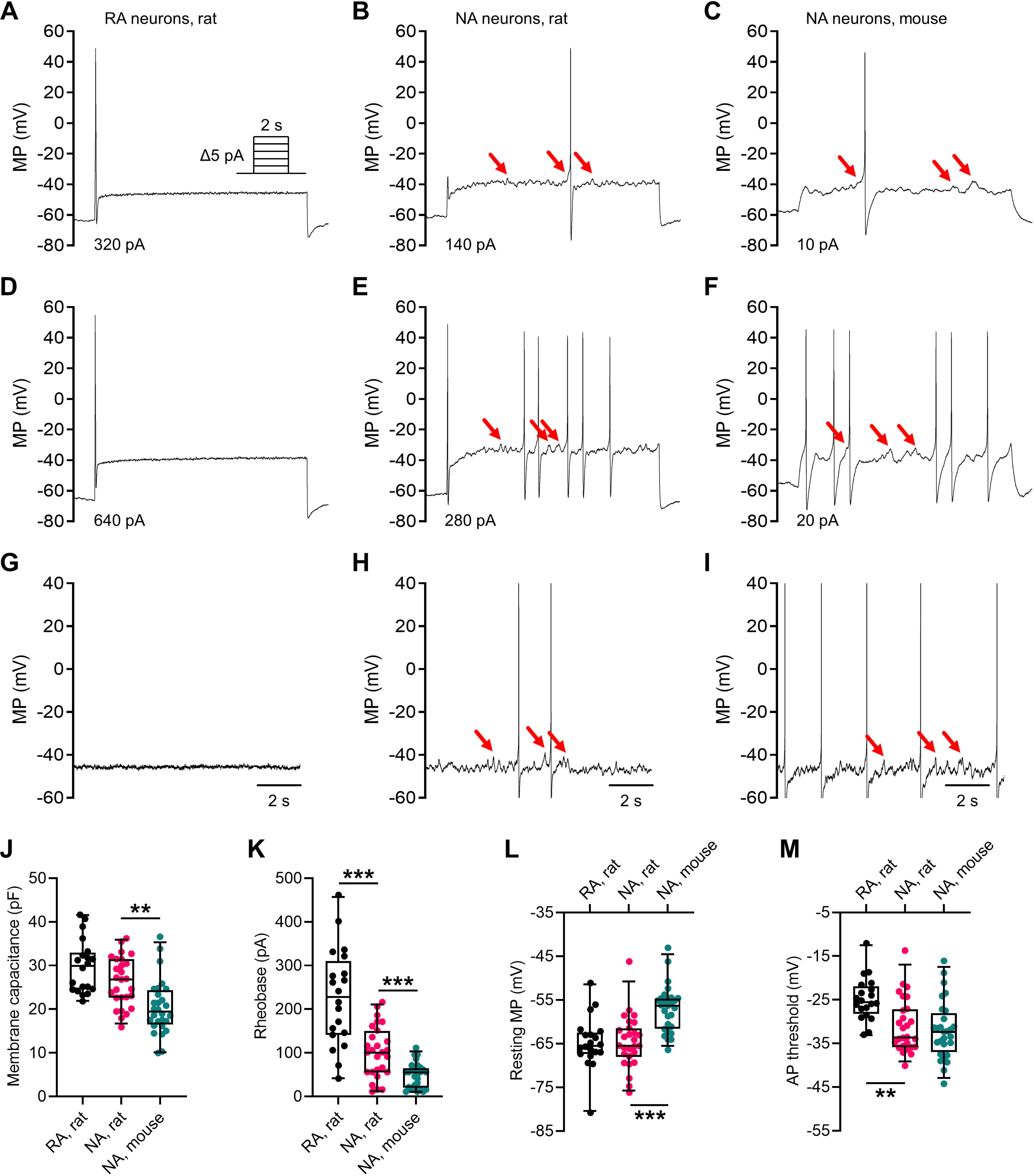
Membrane potential fluctuations, action potentials, and excitability properties in nociceptors dissociated from rodents. Example response patterns of RA and NA neurons at rheobase (A-C) and 2x rheobase (D-F). (A inset) Neurons were incrementally injected with brief pulses of depolarizing current from a −60 mV holding potential. (G-I) Prolonged depolarization to a −45 mV holding potential was used to measure DSFs under comparable conditions across all neurons. Large DSFs marked in NA neurons (red arrows). (J-M) Membrane properties measured in neurons (rat RA n = 20, rat NA n = 27, mouse NA n = 28). The mouse NA neurons exhibited increased excitability compared to the rat NA neurons as indicated by a significantly smaller membrane capacitance (J), lower rheobase (K), and depolarized membrane potential (L). The rat RA neurons were less excitable with higher rheobase values (K) and depolarized AP thresholds (M). Data in (J-M) shown as the median ± 95% CI with all points included. Data in (J) were analyzed using a 1-way ANOVA (F(2, 72) = 15.05, *P* < 0.0001) followed by Dunnett’s multiple comparisons test. Data in (K) were analyzed using a Brown-Forsythe 1-way ANOVA (F(2.00, 30.78) = 31.95, *P* < 0.0001) followed by Dunnett’s T3 multiple comparisons test. Data in (L-M) were analyzed using Kruskal-Wallis tests (KW = 26.12, *P* < 0.0001 and KW = 16.37, *P* = 0.0003, respectively) followed by Dunn’s multiple comparisons test. \*\**P* < 0.01, \*\*\**P* < 0.001. ANOVA, analysis of variance; CI, confidence interval; MP, membrane potential.

Finally, it is assumed that signal analysis techniques like the fast Fourier transform (FFT) capture the dominant frequencies associated with membrane potential oscillations (or fluctuations as we refer to them, to include highly irregular changes in potential) in rodent sensory neurons [20–27]. To test this assumption, a subset of the −45 mV recordings (n = 10 neurons per group) were analyzed using the FFT. The dominant frequencies were highly variable (ranges: rat RA = 0.2-50 Hz, rat NA = 0.15-15 Hz, mouse NA = 0.15-10 Hz; FFT results are available online, see Program and Data Availability section in Methods) and did not completely agree with the published FFT results of 15-107 Hz for NA-like small-diameter neurons isolated from rats [20]. This suggested the FFT may not be suitable for making inferences about the DSFs.

Next, we used FIBSI to isolate the basic dimensional properties of all DSFs in the RA and NA neuron datasets to determine whether DSFs differed between neuron types and rodent species (**Fig 2A-C**). Current-clamp recordings had been performed using the same equipment as reported previously [17,18], so we used published cutoffs (≥1.5 mV and ≥20 ms) to eliminate from analysis low-amplitude signals that are likely noise from equipment. For FIBSI analysis, a running median window of 1 s corresponding to 20,000 samples along the x-axis was used, and no filters or transformations were applied to the raw data prior to signal detection.

**Fig 2.**
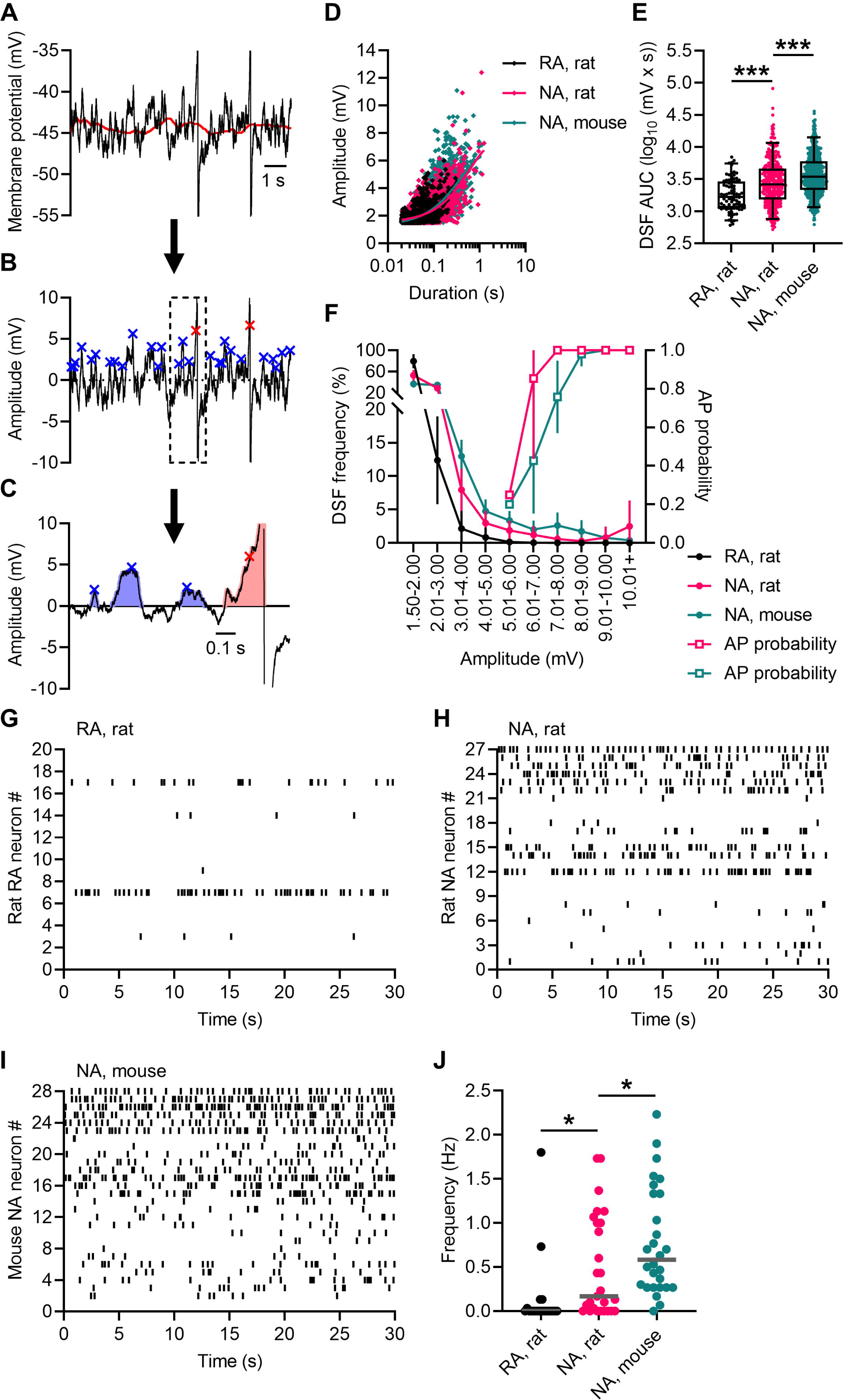
Processing whole-cell current-clamp recordings to measure membrane potential fluctuations in rodent nociceptors. (A-C) Event detection workflow in a whole-cell current-clamp recording measured from a rat NA neuron (APs are truncated). Order of operations: (A) calculate the running median (red line, window = 1 s) based on the raw voltage and time coordinate data; (B) convert the running median to y = 0 and calculate residuals, identify peaks of DSFs (blue X) that meet user-defined cutoffs (≥3.0 mV amplitude, ≥20 ms duration in this example), and manually approximate the amplitude of suprathreshold DSFs (red X) that produce an AP (see Methods for amplitude substitutions); (C) visual confirmation of the DSF dimensional characteristics. Descriptive information for the subthreshold DSFs include start, peak, and end times, amplitude, duration, and AUC (blue highlight) to the converted running median at y = 0. Descriptive information for the AP waveform (pink highlight) includes the aforementioned properties. (D) Scatter plot of DSF dimensions and exponential plateau curves (cutoffs: ≥1.5 mV and ≥20 ms; rat RA n = 20 neurons, rat NA n = 27 neurons, mouse NA n = 28 neurons). Data were fit using exponential plateau models for each neuron type; a single model did not fit all data (F(6, 5689) = 6.696, P < 0.0001). Refer to **S1 Table** for model parameters. (E) The AUC of the DSFs ≥3 mV was significantly larger in rat NA neurons compared to rat RA neurons. The DSFs in Mouse NA neurons were also significantly larger than in rat NA neurons. (F) The NA neurons exhibited rightward shifts in DSF frequency (%) per amplitude bin compared to the RA neurons. In NA neurons, DSFs ≥5 mV triggered APs and the AP probability rapidly increased to 1 at 7-9 mV. Raster plots depicting frequencies of DSFs ≥3 mV in (G) rat RA neurons, (H) rat NA neurons, and (I) mouse NA neurons. The rat and mouse NA neurons with suprathreshold DSFs and ongoing activity at −45 mV correspond to numbers 22-27 and 23-28 on the y-axis of (H-I), respectively. (J) The mean frequencies for DSFs ≥3 mV were significantly higher in the NA neurons. Data in (E) shown as the median ± 95% CI with all points included. The raw AUC data collected for (E) were log transformed for the use of parametric statistics. Transformed results were analyzed using a 1-way ANOVA (F(2, 971) = 38.90, *P* <0.0001) followed by Sidak’s multiple comparisons test. Data in (J) shown as medians (gray line) and analyzed using a Kruskal-Wallis test (KW = 23.85, *P* < 0.0001) followed by Dunn’s multiple comparisons test. \**P* < 0.05. AUC, area under the curve.

To our knowledge, no statistical model has been reported that describes the relationship between DSF amplitude and duration. Such a model would provide insight into the channel opening and closing dynamics driving DSF generation in nociceptors. This model would also provide a reference against which the effects of various pharmaceuticals can be compared and future treatments for pain may be developed. In order to approach this is an unbiased manner, we employed the Akaike information criterion to compare the quality of fit of several different nonlinear regression models to the neuron datasets. Overall, the exponential plateau model was a better fit for the RA and NA neurons compared to the Gompertz, Logistic, and Malthusian models (see **S1 Table** for model parameters). We then asked whether the exponential plateau model could fit all data points using the same set of fitting parameters, but a separate set of parameters was necessary for each neuron type (F(6, 5689) = 6.696, *P* < 0.0001; **Fig 2D**). All models tested, including the exponential plateau model for each neuron type, failed to pass for homoscedasticity (*P* < 0.0001 for each model), and the 95% CI for the RA neuron model could not be calculated by the GraphPad Prism software. These results suggest interpretations based solely on statistical models should be made with caution, but they might provide some insight into the relationship between DSF amplitude and duration.

Making comparisons based solely on DSF amplitude (like done previously [17,18]) limits interpretation of the results because it omits any influence duration may have on DSF size, as the DSFs do not exhibit regular waveforms. Therefore, we asked whether the total AUC of DSFs differed between neuron types. A cutoff of ≥3 mV was chosen because it is at the lower bound of the range of amplitudes that may be functionally relevant under pain-associated conditions when neurons are depolarized [17,18]. The AUC of DSFs in rat NA neurons was larger than in RA neurons, and the AUC of DSFs in mouse NA neurons was the largest (**Fig 2E**). We then binned DSFs by amplitude to determine their frequency of occurrence (reported as a percentage of total DSFs generated in each amplitude bin), and the amplitude necessary to generate APs (**Fig 2F**).

The mean frequency of DSFs per amplitude bin was higher in NA neurons than RA neurons for rats. No DSFs triggered APs in the RA neurons, but a fraction of the large amplitude DSFs (≥5 mV) in NA neurons did trigger APs (**Fig 2F**). Mouse NA neurons tended to require larger DSFs to trigger APs before the AP probability plateaued. Finally, we used raster plots to depict the differences in frequencies of DSFs ≥3 mV between RA and NA neurons (**Fig 2G-I**). Few RA neurons generated DSFs ≥3 mV (**Fig 2G**). The frequency of DSFs ≥3 mV was significantly higher in NA neurons from mice (**Fig 2I**) than rats (**Fig 2H-J**). The frequencies of DSFs ≥3 mV detected in NA neuron recordings by FIBSI were orders of magnitude lower than the oscillation frequencies reported based on similar measurements using the FFT [20].

### 2.2. Using fluorescence contrasted visualization and FIBSI to analyze gut motility in the larval zebrafish

Direct intestinal injections and oral gavage can be used to deliver dyes into zebrafish larvae [28,29]. However, these methods are invasive, can be deterrents without adequate technical expertise, and may directly influence motility. We first determined whether immersing awake larvae in embryo medium (E3) media supplemented with Nile Red dye – without using an egg emulsion to promote feeding [30] – was a viable non-invasive alternative for staining the intestinal luminal space (**Fig 3A-B**). No acute or long-term (≥24 hours) effects of exposure on general activity levels and behavior, and no mortality were observed at the Nile Red dose used (0.01 μg/mL). Video playback of the fluorescence time-lapses (**S2 Movies**) confirmed contractions propagated at different rates in the different regions of the gut, and in opposing directions (i.e., retrograde and anterograde [31]) from a site of pacemaker-like activity near the end of the foregut [32]. A total of 19 ROIs were evenly positioned along the foregut, midgut, and hindgut regions of the intestinal tract (**Fig 3B-C**). We predicted that the Nile Red fluorescence intensity would change periodically, reflecting contraction waves traveling through the gut (**Fig 3D-E**); retrograde contractions coincide with a decrease in intensity in the foregut, anterograde contractions coincide with an increase in intensity in the midgut and hindgut, and rapid movement of the cloaca coincide with high-frequency oscillations in intensity in the hindgut. Indeed, contractions were visible in the fluorescence intensity recordings as they propagated rhythmically through the intestinal tract at <0.1 Hz, and the directional changes in intensity matched our predictions (**Fig 3F**).

**Fig 3.**
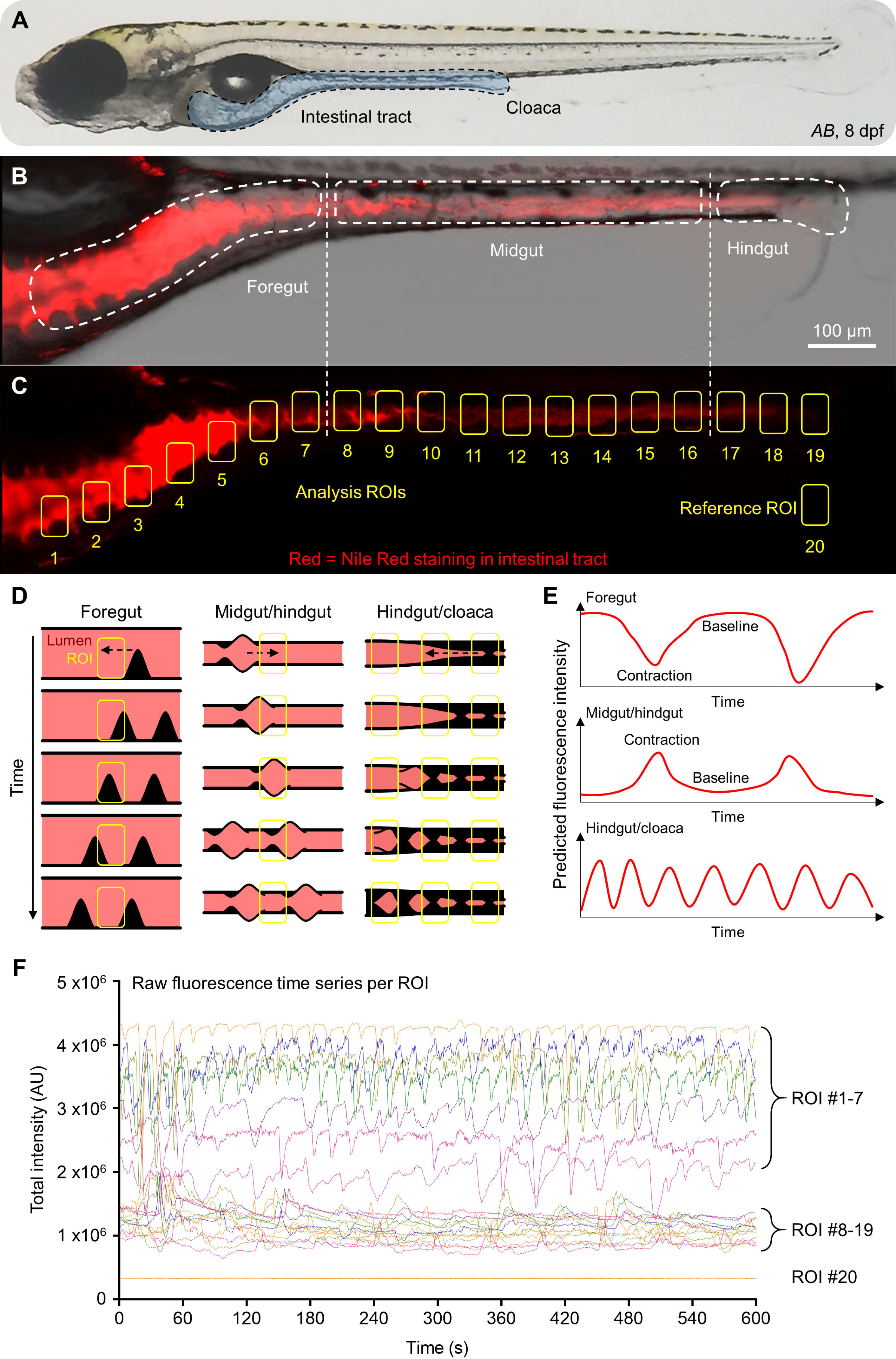
Fluorescent-contrasted visualization of gut motility in larval zebrafish. (A) Wild-type *AB* zebrafish larvae at 8 dpf. The intestinal tract is outlined and highlighted in blue. (B) Merged bright field and red fluorescence (Nile Red dye) image with different functional segments of the intestinal tract labeled. (C) Fluorescence image of (B) depicting positions of ROIs used to measure changes in fluorescence intensity resulting from muscle contractions. (D-E) Models and predicted time series depicting fluorescent fluctuations from baseline in ROIs during propagation of retrograde contractions in the foregut and anterograde contractions in the midgut/hindgut. The high-frequency movements of the cloaca were predicted to generate oscillatory changes in fluorescence. (F) Time course of raw fluorescence intensity changes measured from 20 ROIs in (C) in a single larva (traces color coded for visual differentiation). dpf, days post-fertilization.

During our development process, we noticed that some recording periods were primarily high fluorescence, with contractions signaled by a sudden reduction in fluorescence (a trough event), whereas other recordings were primarily low fluorescence, punctuated by sudden increases in fluorescence (a peak event). This is likely related to differing contraction physiology in the foregut, midgut, and hindgut. In order to increase the comparability of these events and standardize the identification of amplitude, we implemented a second normalization procedure that traces a fitted line peak-to-peak (or trough-to-trough) of events identified in the first round of normalization (i.e., sliding median). This new line is used as the reference against which the data is compared and events identified again. The peak-to-peak function was applied to the foregut (ROIs 1-7) and hindgut (ROIs 17-19), and the trough-to-trough function was applied to the midgut (ROIs 8-16). The following running median window sizes were used: 50 x-samples for ROIs 1-7 and 10 x-samples for ROIs 8-19.

Processed fluorescence intensity data for the foregut (**Fig 4A-C**), midgut (**Fig 4D-F**), and hindgut ROIs (**Fig 4G-I**) revealed several features: foregut contractions were overall larger in amplitude, foregut and midgut contractions had similar durations, and the high-frequency oscillations coinciding with cloaca movement in the hindgut were overall shorter and smaller. These features mirrored observations made during video playback and predictions about signal shape (**Fig 3D-E**). A cutoff of ≥0.05 amplitude (F/F0 units) and duration of ≥10 s (duration cutoff inferred from published contraction frequencies and intervals, see [31,33–35]) was used to isolate low-frequency muscle contractions (quadrant 4, Q4) in ROIs 1-19 from noise (Q1-3) Here, noise refers to signal artifacts and movements of the gut not related to the muscle contractions of interest observed in the videos. A separate cutoff of ≥0.05 amplitude and duration window of 5 s ≤ x ≤ 10 s was used to isolate the rapid movements of the cloaca present in ROIs 17-19. Raster plots for Q1-3 (**S3 Figure**) did not depict rhythmic activity indicative of peristalsis in either unfed or fed larvae. In contrast, raster plots for Q4 in unfed (**Fig 4J**) and fed larvae (**Fig 4K**) exhibited key characteristics of peristalsis also observed in spatiotemporal maps [31]: anterograde (purple arrows) and retrograde (green arrows) waves traveled rhythmically through the intestinal tract. These waves originated from a pacemaker-like region near the end of the foregut (blue highlight) [31,32].

**Fig 4.**
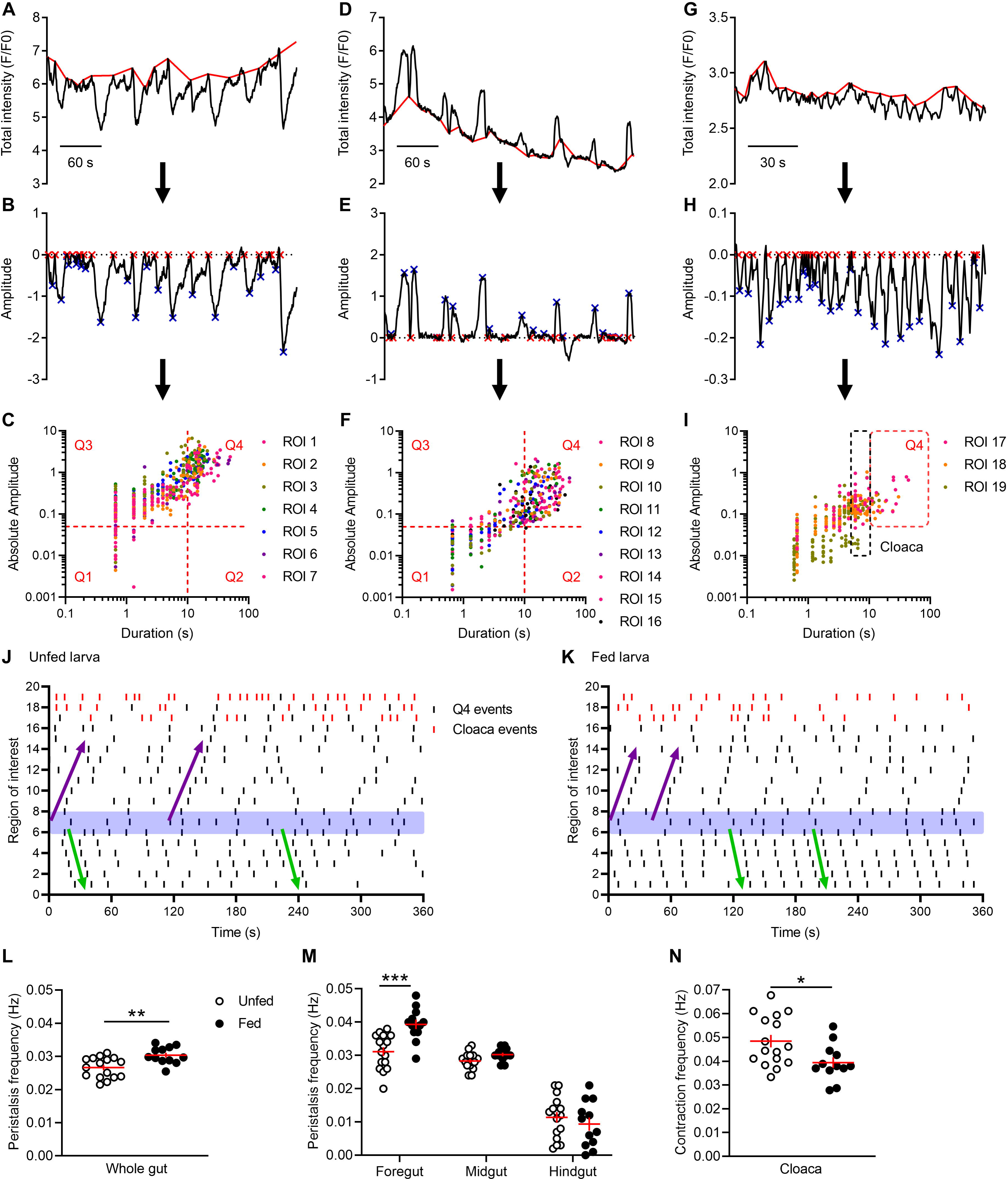
Processing fluorescence time series datasets using FIBSI to analyze gut motility in larval zebrafish. Event detection workflow for fluorescence time series corresponding to ROIs #1-7 in the foregut (A-C), #8-6 in the midgut (D-F), and #17-19 in the hindgut (G-I). The residuals were calculated from the peak-to-peak or trough-to-trough trend line (red) and the trend line was normalized to y = 0. Event peaks (blue X) were identified only for the waveforms with start/end times that crossed y = 0 (red X). After processing, event durations and amplitudes were plotted for inspection and filtering (1 dot = 1 event, events are color coded by ROI). Cutoffs (dotted red lines, amplitude of ≥0.05 and duration ≥10 s) were applied to isolate low-frequency contractions in Q4 of al 19 ROIs. Oscillatory movement of the cloaca was isolated in ROIs 17-19 using a cutoff window (dotted black line, amplitude cutoff of ≥0.05 amplitude and 5 s ≤ x ≤ 10 s duration). Raster plots of events in Q4 and cloaca in unfed (J) and fed (K) larvae depict low-frequency retrograde (green lines) and anterograde (purple lines) contractions traveling through the gut. Contractions originated from a region of pacemaker-like activity near the end of the foregut (blue shading at ROIs #6-8). (L) Mean contraction frequencies across the length of the whole gut (ROIs #1-19) were significantly increased in fed (black circles) compared to unfed (white circles) larvae. (M) Fed larvae had significantly increased mean contraction frequencies in the foregut (ROIs #1-7), but exhibited no differences from unfed larvae along the midgut (ROIs #8-16) and hindgut (ROIs #17-19). (N) Mean contraction frequencies for the cloaca were significantly decreased in fed larvae. Points in (L-N) represent the mean per unfed (n = 16) and fed (n = 13) larvae and shown as the group mean ± SEM. The initial comparison between groups in (L) was made using an unpaired *t* test and gut region-specific comparisons in (M) were made using a 2-way ANOVA (Group factor F(1, 78) = 6.116, *P* = 0.0156; Region factor F(2, 78) = 188.6, *P* < 0.0001; Interaction F(2, 78) = 7.460, *P* = 0.0011) followed by Sidak’s post-hoc test. Data in (N) were compared using an unpaired *t* test. \**P* < 0.05, \*\**P* < 0.01, \*\*\**P* < 0.001. SEM, standard error of the mean.

We then asked whether the spatiotemporal information extracted from our time-lapse datasets using FIBSI could be used to detect differences in gut motility between unfed and fed larvae. A recent study using image velocimetry and spectral analysis reports differences in motility between unfed and fed zebrafish larvae at 7 dpf [35], but the study is limited to the midgut, and no other comparable datasets have been published. First, we confirmed that muscle contraction frequencies did not differ between unfed *AB* (0.026 ± 0.001 Hz, n = 8) and unfed transgenic *Tg(−8.3phox2bb:Kaede)* larvae (0.028 ± 0.001 Hz, n = 8; unpaired *t* test *P* = 0.158). Contraction frequencies in the cloaca also did not differ between the two lines (0.046 ± 0.004 Hz and 0.051 ± 0.003 Hz, respectively; unpaired *t* test *P* = 0.407). Therefore, the two lines were pooled to increase statistical power for comparison to fed *AB* larvae. Muscle contraction frequencies were higher in fed compared to unfed larvae (**Fig 4L**).

We then asked whether motility in fed larvae was higher along the entire gut, or if the increased motility was limited to a specific functional gut segment. The frequency of retrograde contractions in the foregut was higher in fed compared to unfed larvae (**Fig 4M**). In contrast to the prior study [35], we observed no differences between unfed and fed groups for anterograde contractions in the midgut or hindgut (**Fig 4M**). Finally, we observed a significant decrease in the frequency of cloaca movements in fed compared to unfed larvae (**Fig 4N**). The data collected using our fluorescence contrast-based assay of gut motility, in conjunction with signal detection and data analysis using FIBSI provided novel insights into individual functional gut segments of larval zebrafish. We were able to characterize peristalsis along the whole intestinal tract as well as high-frequency, low-amplitude oscillatory movements of the cloaca.

## 3. Discussion

### 3.1. Overview

In this study we introduce a new signal detection algorithm implemented in the free program FIBSI that can be used to analyze a diverse array of non-linear, dynamic and stochastic experimental datasets by extracting biologically meaningful signals from background noise. As experimental biologists, we wanted to develop FIBSI to be particularly suited to the needs of other experimentalists. As such, strengths of FIBSI include its capability to process non-ideal datasets such as those we included here to demonstrate the utility of the program. These may include datasets with low signal-to-noise ratios, irregular signal shapes, and low-resolution (i.e., under-sampled) time series. In contrast to many other signal detection programs, FIBSI requires no modification of the raw data, and minimal ad-hoc assumptions regarding data structure and signal shape are needed prior to processing and signal detection. That is, FIBSI captures quantitative information for all signals contained with the dataset. We validated the performance of FIBSI using two different types of measurements: electrophysiological whole-cell current-clamp recordings from rodent sensory neurons and fluorescence intensity imaging data capturing muscle contractions in the larval zebrafish intestinal tract. Hypothesis testing based on prior biological knowledge – two prior pain-related rodent studies [17,18] and established methods for measuring intestinal motility in vertebrates [31–40] – was used to draw novel conclusions about the analyzed datasets. The results of these tests are summarized and discussed below.

### 3.2 Membrane potential fluctuations differ between nociceptors in two rodent species

We were the first to automate the quantification of DSFs and demonstrate their functional importance for AP generation in putative C-type nociceptors using the prototype algorithm upon which FIBSI is based [17]. We showed that DSFs bridge the gap between the resting membrane potential and AP threshold of a neuron and presumably play a role in driving ongoing pain-related information from nociceptors [41,42]. We have also begun to identify associated cellular mechanisms (e.g., dependence upon cell signals such as cyclic AMP-dependent protein kinase and exchange protein activated by cyclic AMP-associated pathways) [17,18,43] to better understand how DSFs are generated and under what conditions they can be inhibited. Others have also noted their potential role in driving other pain-related conditions in rodents [44] and in humans [45]. Given the limited research on DSFs, our current study is the first to make direct comparisons between similar nociceptors in two rodent species commonly used in pain-related studies. This was done to 1) confirm whether nociceptors in rats and mice function similarly, and 2) to validate FIBSI in its new, generalized form using the same data type as in our original studies [17,18].

The NA neurons isolated from rats and mice recapitulated several of the specializations we have described that promote ongoing activity and ongoing pain [17,18,41–43]. Under naïve conditions (i.e., no injury), rat NA neurons are clearly more excitable than RA neurons. A novel finding presented here is that mouse NA neurons are more excitable than rat NA neurons with enhanced rheobase values and depolarized resting membrane potentials. A plausible explanation for the species-specific differences in NA neuron excitability is that the neurons in mice may be smaller in size, as suggested by the smaller membrane capacitance measures. This raises the interesting question of whether the membrane densities and compositions of the channels underlying the DSFs differ between species; this is of ongoing interest in the Walters lab and under investigation. These specializations are expected to prime the NA neurons to activate more readily in response to intra- and extracellular signals associated with nociceptive and potentially painful stimuli. This hypothesis is further supported by our analysis of DSFs using FIBSI. The mouse NA neurons generate larger, more frequent DSFs than rat NA neurons. However, the greater DSF size and frequency, and necessity for larger-amplitude DSFs to generate APs in mice compared to rats is unexpected because naïve rats and mice have similar AP voltage thresholds while the mice have a more depolarized RMP compared to rats. We also compared exponential growth models to describe the dimensional properties of the DSFs and found different models were necessary for each neuron type. These models need to be compared further under other pharmacological and pain-related conditions in order to better understand the differences between species. Finally, the frequency results we obtained in our FFT analysis did not recapitulate the results for the same type of neuron [20], and other studies do not fully report their results for comparison [21–27]. One possible explanation for the discrepancy is that the longer sampling duration used (30 s compared to 1-4 s in the original study [20]) gave a more representative image of the fluctuations and activity in each neuron, which may influence the measured frequencies. Using FIBSI, frequencies for DSFs ≥3 mV in NA neurons depolarized to −45 mV peaked at 1-2 Hz. This finding is similar to our spinal cord injury and serotonin-induced inflammatory conditions, under which frequencies for DSFs ≥5 mV peak at ~1.5 Hz [17]. Given these similarities, it is unlikely we would observe a shift in the FFT signal to 20-107 Hz for NA neurons depolarized to −45 mV, even if we used the same injury condition as the prior study (sciatic nerve injury [20]). Taken together, these observations suggest the FFT is not an effective tool for analyzing highly irregular DSFs in membrane potential, as opposed to the sinusoidal oscillations examined primarily in large DRG neurons previously (most of which are not nociceptors) [20–27]. Unlike the FFT, the descriptive data generated by FIBSI can be concretely linked to precise DSFs of interest and dissociated from other biological and non-biological signals in the current-clamp recordings.

Our findings suggest two assumptions that should be avoided when investigating DSF mechanisms: 1) DSFs of one species are indistinguishable from DSFs of another, even in what appear to be functionally equivalent nociceptors, and 2) DSFs in their measured form represent unitary events. Future studies can test whether DSFs usually represent summation of discrete events and if underlying unitary components can be identified based on waveform characteristics (see [46,47]). The use of FIBSI and similar analytical approaches used to study DSFs will improve our understanding of basic nociceptor physiology in multiple species and help to identify fundamental mechanisms for driving pain that may be shared by rodents and humans.

### 3.3 Validation of a simple, fast, and cost-effective assay of gut motility in larval zebrafish

The current approach for assessing gut motility in vertebrates *in vivo* and *ex vivo* is to generate spatiotemporal maps of the intestinal tract and measure changes in pixel opaqueness as indices of movement [31,32,34,36–40]. Inspection of these maps can be done manually but the process is laborious and possibly subject to human unconscious bias. More recent studies have improved the analysis of gut motility in that the process has been semi-automated, which also enables assessment of a wider range of parameters (e.g., contraction amplitude in μm). However, these methods require custom microscopy equipment [48], complex image velocimetry and spectral analysis [35], and/or use of customized “in-house” [49] or proprietary software [33,40]. Here, we developed a simpler, faster, and more cost-effective assay using fluorescence contrast-based imaging to visualize gut motility *in vivo*. The resulting time series datasets can be processed using FIBSI to analyze muscle contractions. While 8- and 16-bit grayscale bright field images can also be analyzed using FIBSI, fluorescence intensity measurements have a larger dynamic range, and the resulting time series have a higher signal-to-noise ratio (i.e., contractions are more visible) The advantage of FIBSI over existing programs used to analyze gut motility is its user-friendliness and free availability.

Our attempts to validate the fluorescence contrast-based gut motility assay and the performance of FIBSI against prior studies was challenging due to the fact that many reports do not include sufficient experimental details (e.g., feeding regimen, gut region imaged, and methodology used to calculate contraction frequencies). It is important that future studies are explicit in their reporting to facilitate more direct comparisons between studies and method development. Some studies measure motility in only select regions of the gut [29,35,50] while others measure motility throughout the whole gut and pool the results [33]. We contend calculating the frequency of retrograde and anterograde contractions separately [31,34] gives a more accurate representation of gut function and how its luminal contents may be processed mechanically. Our gut motility assay and FIBSI captured important characteristic features of peristalsis also observed in the spatiotemporal maps: anterograde and retrograde contractions originated from a site of pacemaker-like activity posterior to the intestinal bulb [31,32] near our delineation of the foregut and midgut. Observation of these features increased our confidence in the results. The contraction frequencies obtained using FIBSI recapitulated the published range of frequencies for unfed and fed larvae [33,34,50]. The only published study to directly compare gut motility in unfed and fed zebrafish larvae used measurements from the midgut, and it reports feeding at 5-6 dpf increases motility at 7 dpf from 0.035 Hz to 0.041 Hz [35]. We observed similar rates (~0.03 Hz) in the midgut, but feeding zebrafish live paramecia 1 day before testing did not increase the rate of anterograde contractions in the midgut. Instead, we found that feeding increased retrograde contractions in the foregut. Another novel observation was that feeding decreased the rate of oscillatory movements of the cloaca. Adult shorthorn sculpin – another teleost species – respond to feeding in a similar manner; orally-directed (retrograde) standing contractions increase in frequency while slow, anally-propagating (anterograde) contractions occur less frequently [49]. We did not observe a decrease in anterograde contractions, but it is likely that the increase in retrograde contractions in sculpin and zebrafish serve a similar function to displace and propel large quantities of luminal contents [49]. In zebrafish larvae, the oscillatory motions of the cloaca may constitute sphincter-like contractions that control the timing of defecation by keeping the luminal contents inside the fish rectum. Evidence suggests the caudal segments of the adult zebrafish intestine may share functional similarities with the human rectum [51], but to our knowledge there are no published reports of sphincter-like function in the hindgut of larval zebrafish. It is possible that a reduction in the oscillations (i.e., relaxation) may facilitate more frequent removal of waste products, but this has yet to be confirmed using transit experiments. Our findings suggest that larval zebrafish are capable of generating functionally complex gut motility patterns in response to feeding, possibly similar to other adult teleost species. This adds to their tractability as a powerful model organism for *in vivo* imaging. The significance of these gut region-specific phenomena is becoming increasingly apparent (for review [52]), but these complex motility patterns require further attention using methods that yield functional information of the whole gut. Future improvements to our motility assay and FIBSI could link changes in contraction amplitudes to changes in the luminal dimensions of the intestinal tract, similar to using image velocimetry to track movement [35].

There is a pressing need for improved imaging and analysis techniques that can be used to address gut motility in altered physiological states and disease [53]. Analysis with FIBSI yields experimentally valid and reproducible results, and it should be useful to researchers measuring gut motility in zebrafish as a functional output of the enteric nervous system (ENS). The ENS regulates multiple gastrointestinal functions (e.g., gut motility, hormone secretion). Deficiencies in the development and function of the ENS can lead to debilitating neurological disorders (e.g., Hirschsprung’s disease) [54–58]. Other factors of interest associated with ENS function (e.g., stress, microbiome, inflammation, visceral pain) [59–61] have traditionally been tested using rodent models. Recent reviews have highlighted the validity of using zebrafish as an analogous model to address some of the above questions [53,62–65], and the application of our motility assay and FIBSI to this system will facilitate such research.

### 3.4. Current and future applications of FIBSI

We originally designed the prototype version of FIBSI (first report [17]) for the singular purpose of quantifying DSFs and APs in nociceptors and incorporating the methodologies of others (e.g., estimating AP threshold [66]) within the field. However, collaborative efforts led us to realize the broader applicability of the program to other researchers and types of datasets. Additional features (e.g., filtering methods, down-sampling, trimming) are included for those interested, and we encourage others to make modifications to the existing code (see availability section in Methods) and/or insert their own functions. The signal detection algorithm was tested solely on datasets with fixed sampling rates. Although the minimum requirement for identifying an event is two consecutive x values with the same sign, it is possible some assumptions we have made in other aspects of the algorithm may lead to inaccurate identification of events in datasets with variable sampling rates. Testing and implementing different methods of data interpolation could alleviate this limitation but has not been done by our research group.

We speculate that FIBSI could theoretically work with almost any type of x,y dataset where the measured parameter exhibits deviations from baseline, as FIBSI is adapted to account for changes in the baseline level of activity via the running fit function. This is an improvement over arbitrary decisions for baseline or rudimentary measures (e.g., simple average) that assume the system being studied is fixed. Although we have only fully tested FIBSI for processing of voltage fluctuation time series and fluorescence intensity time series, ongoing projects are currently adapting FIBSI to process other types of datasets (e.g., *in vivo* nerve recordings and calcium imaging in rodent and zebrafish neurons), and to investigate physiological states and disease models pertinent to our research groups. Other examples of biological datasets that may be processed and analyzed using FIBSI include animal vibrations (e.g., honeybee waggle dance [67]) and sounds (e.g., ultrasonic vocalizations [68,69]). In summary, there exists broad potential for FIBSI to facilitate signal analysis of diverse types of raw data by biologists having quite different backgrounds in quantitative analysis.

## 4. Methods

### 4.1. Description of frequency-independent biological signal identification (FIBSI) program

The signal detection algorithm used by FIBSI was generalized from our initial study where we identified DSFs in nociceptors [17]. It is an extension of the Ramer-Douglas-Peucker algorithm for identifying significant shapes that ought to be retained in an image while reducing the total number of points needed to represent said image [15,16]. The FIBSI program was developed and tested on Ubuntu Linux and Windows 10 operating systems. The data processing is as follows:

- A vector *Y*_*trace*_ in which *t* = time points and *y* = independent measured variable in an x, y series is received as input

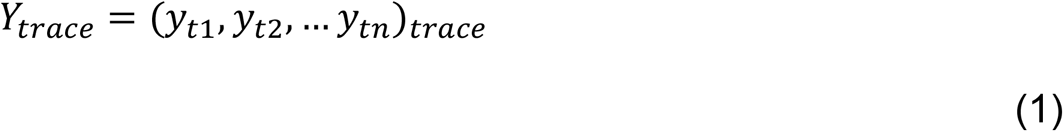
- The vector can be normalized to an external reference if selected

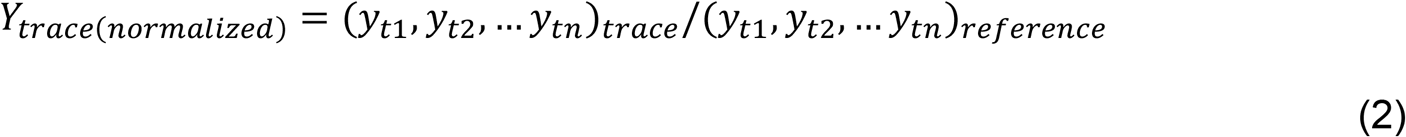
- A fitted line is generated where *f* = function used (e.g., running median, least-squares linear regression)

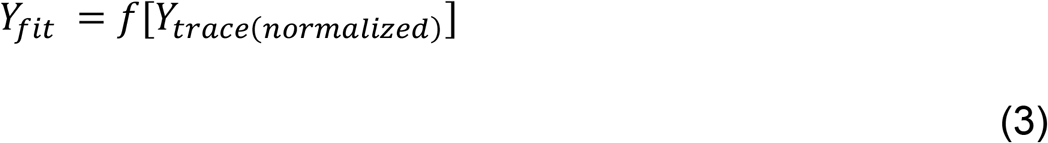
- The residuals to the fitted line at each given point are calculated

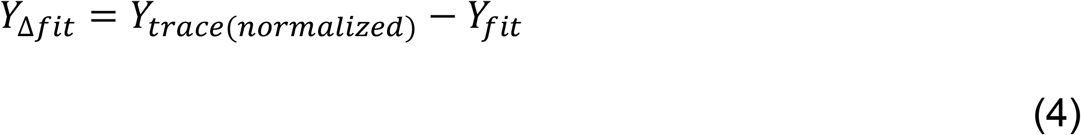
- Signal/event detection is performed by dividing *Y*_Δ*fit*_ into discrete local maxima ‘above’ and local minima ‘below’ waveforms in relation to *Y*_*fit*_:

a. We first define that *y* values indexed at time points where *y* ≥ 0 are elements of ‘above’ time points, and *y* values at time points where *y* < 0 are elements of ‘below’ time points:

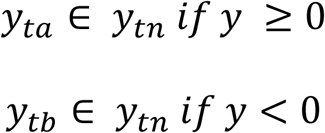
b. We then define the ‘above’ and ‘below’ vectors in *Y*_Δ*fit*_ where *m* is the length of the ‘above’ vector:

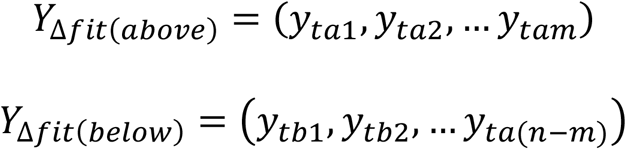
c. We then define the set of criteria for discrete ‘above’ events, where *y* = *f*(*T*), and *t*_*i*_ ∈ *T* where *t*_*i*_ is a local maxima if the following conditions are met:

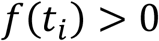 Start (rise) time: *t*_*R*_ ∈ *T* such that *t*_*R*_ < *t*_*i*_ and *f*(*t*_*R*_) = 0 End (fall) time: *t*_*F*_ ∈ *T* such that *t*_*F*_ > *t*_*i*_ and *f*(*t*_*F*_) = 0 Components of the rising phase: *R* = {*t* ∈ *T* | 0 < *t*_*R*_ < *t*_*i*_} Components of the falling phase: *F* = {*t* ∈ *T* | *t*_*i*_ < *t*_*F*_ < *t*_*n*_} R must not contain numbers that cross zero: (∀_*t*_ ∈ *R*)*f*(*t*) > 0 F must not contain numbers that cross zero: (∀_*t*_ ∈ *F*)*f*(*t*) > 0 An ‘above’ event, *Z*, is the union of components in *R* and *F*: *Z* = *R* ∪ *F* (∀_*t*_ ∈ *Z*)∄_*t*_ where *f*(*t*) > *f*(*t*_*i*_) ⇒ *t*_*i*_ is the time of the local maximum, *t*_*R*_ is the start time, and *t*_*F*_ is the end time ∴ *Z* is the set of numbers contained within a single ‘above’ event
d. The local minima for a ‘below’ event is therefore defined as *f*(*t*_*i*_) < 0 and a similar set of rules:

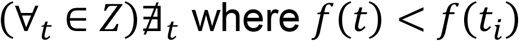
- Events can be excluded via user-defined duration and amplitude cutoffs, and replaced with the line of best fit connecting the start and end points of the excluded event; this is a generalization of the AP exclusion method described previously [17]
- Refitting can be applied; this allows additional use of a trend line calculated by tracing the peak-to-peak pathway of identified ‘above’ or ‘below’ events; steps (3-4) are repeated to generate a new *Y*_*fit(peak)*_ and *Y*_Δ*fit(peak)*_; troughs can also be used to generate *Y*_*fit*(*trough)*_ and *Y*_Δ*fit(trough)*_
- If exclusions and/or refitting methods are applied, then event detection can be performed again
- Event directionality (above or below), dimensions (duration, amplitude, AUC), and indices (start, peak, end) are tabulated; graphs and comma-separated x, y series can be generated as optional outputs

### 4.2. Data analysis

Analysis of raw electrophysiological and fluorescence time series data was performed with FIBSI written using the Anaconda v2019.7.0.0 (Anaconda, Inc, Austin, TX) distribution of Python v3.5.2, and the NumPy and matplotlib.pyplot libraries. Custom scripts were also written to analyze the electrophysiological data using the FFT. Statistical analyses of the DSF and gut motility data generated by FIBSI were performed using Prism v8.2.1 (GraphPad Software, Inc, La Jolla, CA). Decisions to compare the DSFs in NA neurons between rats and mice, and to compare the DSFs between the NA and RA neurons in rats were made *a priori*. Amplitude substitutions for suprathreshold DSFs were calculated manually as described previously [17,18] (see also the FIBSI tutorial included online). One zebrafish (fed) was omitted from analysis due to a lack of muscle movements within the gut, but blood flow was still observed. This was confirmed during video playback and inspection of the raw fluorescence intensity time series. Statistical significance was set at *P* < 0.05 and all reported *P* values are two-tailed.

### 4.3. Program and data availability

The FIBSI source code, experimental data used in this study, and a tutorial for using FIBSI are available on a GitHub repository titled “FIBSI Program” by user rmcassidy (https://github.com/rmcassidy/FIBSI_program). This has been done to promote the free use, modification, and distribution of FIBSI as a useful tool for researchers. Forking the repository is encouraged and changes will, to the extent possible, be pulled into the original repository as an update.

### 4.4. Description of sensory neuron electrophysiological data used for post-hoc analysis

A subset of electrophysiological data obtained from dorsal root ganglia (DRG) sensory neurons dissociated from 16 male Sprague-Dawley rats and 1 male and 4 female C57BL/6 wild-type mice were used for post-hoc analysis. These rodents were used as naïve controls in prior studies performed by our research group, and detailed methods have been published [18,19]. Rodent experiments were performed in accordance with the International Association for the Study of Pain, the Guide for the Care and Use of Laboratory Animals, and were approved by the Institutional Animal Welfare Committee of the University of Texas Health Science Center at Houston, protocol numbers AWC-18-0035, AWC-15-0122, AWC-18-0134, and AWC-15-0160. Briefly, the DRG were harvested below thoracic vertebrate levels T9 (mice) and T10 (rats) and sensory neurons were enzymatically dissociated for overnight incubation at 37 °C with 5% CO_2_. After 18-24 hours, whole-cell current-clamp recordings were performed. To consistently analyze the DSFs in different neurons and rodent species, only recordings during which neurons were injected with the current necessary to hold the membrane potential at −45 mV for 30 seconds to model pain-associated conditions [17,18] were used. Neurons were sampled at 20 kHz with a 10 kHz Bessel filter, and the time and voltage coordinate data were extracted to comma separated value (.csv) files using PatchMaster v2x90.1 software (HEKA Elektronik, Holliston, MA).

### 4.5. Zebrafish handling and care

All zebrafish care, breeding, and experiments were performed in accordance with the Guide for the Care and Use of Laboratory Animals and were approved by the Institutional Animal Welfare Committee of the University of Texas Health Science Center at Houston, protocol number AWC-19-0078. Adult *AB* wild-type zebrafish (*Danio rerio*) were housed inside the Center for Laboratory Animal Medicine and Care specific pathogen-free aquatic facility in recirculating tanks maintained on a 14 h/10 h light/dark cycle at pH 7.2-7.4 and 26 °C. Zebrafish eggs were obtained following natural spawning; breeding groups (4:3 female to male ratio) were placed into separate breeding tanks the night before spawning. Eggs were first rinsed with sterile E3 (prepared in double distilled water with 10 mM HEPES, 5 mM NaCl, 0.17 mM KCl, 0.40 mM CaCl_2_, and 0.67 mM MgSO_4_, pH 7.2) and then raised in petri dishes filled with E3 supplemented with 0.033 mg/mL methylene blue and 0.013 mg/mL 1-phenyl-2-thiourea to prevent melanin synthesis. The eggs were incubated at 28.5 °C in a diurnal incubator (14 h/10 h light/dark cycle) and E3 was exchanged daily until all embryos hatched, then E3 was exchanged every other day. For feeding, a subset of larvae were incubated with the live prey *Paramecium caudatum* for 2 h at 6 days dpf [70]. The remaining larvae were not fed. In some experiments involving unfed larvae, transgenic *Tg(−8.3phox2bb:Kaede)* zebrafish larvae [71,72] were used in place of wild-type *AB* larvae. Adult transgenic zebrafish were housed and spawned in accordance with IACUC-approved protocol 1144724 by the Uribe lab at Rice University. Fertilized embryos were delivered to the Krachler lab at 0 dpf for rearing. Transgenic embryos were handled the same as wild-type *AB* larvae.

### 4.6. Using fluorescence microscopy to image gut motility in larval zebrafish

At 7 dpf, zebrafish larvae were transferred to 25 cm^2^ (T25) culture flasks (10-20 larvae per flask) filled with 15 mL of E3 supplemented with 0.01 μg/mL of Nile Red (Sigma Aldrich, St. Louis, MO). Flasks were incubated at 28.5 °C. Following incubation (2-4 h), larvae were anesthetized with 0.16 mg/mL tricaine (prepared in double distilled water and buffered to pH 7.2 using 0.5 M PIPES and 1 M NaOH; Sigma Aldrich) and sorted for uptake of Nile Red in the intestinal tract. Larvae with intestinal tracts clearly outlined with Nile Red dye were embedded in 1% low melting point agarose (Promega, Madison, WI) in E3 supplemented with 0.16 mg/mL tricaine in 6-well glass bottom plates (#1.5 high performance cover glass, 0.17±0.005 mm; Cellvis, Mountainview, CA). The intestinal tracts of the larvae were imaged for 6-10 min at 22 °C using an inverted confocal laser scanning microscope (Olympus FluoView FV3000; Olympus Corporation of the Americas Headquarters, Center Valley, PA) using the following settings: 10X objective lens with x1.0 zoom, 1024 × 768 × 1 pixel resolution (1.406 μm/pixel), 1.25-2.5 frames/s sampling rate, 561 nm excitation laser (Galvano) set to 0.5% transmission and 650 V, and a 570-670 nm emission detection window. Post-acquisition, a total of 19 analysis ROIs (fixed position) were drawn over the length of the intestinal tract to measure changes in fluorescence during gut motility, and 1 reference ROI was drawn outside the intestinal tract to correct for background fluorescence (see Results for ROI positioning). Care was taken to use the same ROI dimensions and to position the ROIs consistently between larvae (ROI center-to-center distance = ~55 μm). The Nile Red dye was used as a contrast agent to outline the intestinal tract [30]. Changes in fluorescence during gut motility were due to muscle contractions widening, closing, and moving the intestinal tract. Total fluorescence intensity and time coordinate data per ROI were extracted to comma separated value (.csv) files using Olympus FV31S-SW v2.3.2.169 software. All larvae were euthanized at the end of experiments at 7-8 dpf using 1.6 mg/mL buffered tricaine.

## Acknowledgements

The authors would like to thank members of the Uribe, Krachler, and Walters labs for technical assistance.

## Supporting Information Captions

**S1 table. Parameters for DSF exponential growth models.**

* Denotes probability the model is correct in comparison to the exponential plateau. AICc, Akaike’s criterion; df, degrees of freedom; k, rate constant; YM, max y value; Y0, starting y value.

**S2 movies. Fluorescence contrasted contractions during gut motility in unfed and fed zebrafish larvae.**

(A) Unfed larva corresponding to Fig 3B-C; 1 frame = 0.653 s. (B-C) Unfed larvae; 1 frame = 0.659 s. (D-E) Fed larvae; 1 frame = 0.592 s. Playback set to 30 frames/s for all movies.

**S3 figure. Events detected in quadrants 1-3 for unfed and fed zebrafish larvae do not exhibit peristalsis rhythmicity.**

(A-C) Raster plots of Q1-3 corresponding to the unfed larva depicted in Fig 4J. (D) Time course of fluorescence intensity changes measured from 20 ROIs corresponding to the fed larva depicted in Fig 4K (traces color coded for visual differentiation). (E) Data corresponding to (D) post-processing with FIBSI. The same cutoffs (dotted red lines, duration ≥10 s and amplitude ≥0.05) were applied to distinguish between background noise (Q1-3) and low-frequency contractions. Events are color coded according to their ROI of origin in (D). (F-H) Raster plots for Q1-3 corresponding to the data in (E).

