## Supplementary figures and images for "Frequency-independent biological signal identification (FIBSI): A free program that simplifies intensive analysis of non-stationary time series data"

### Supplemental Figure 3

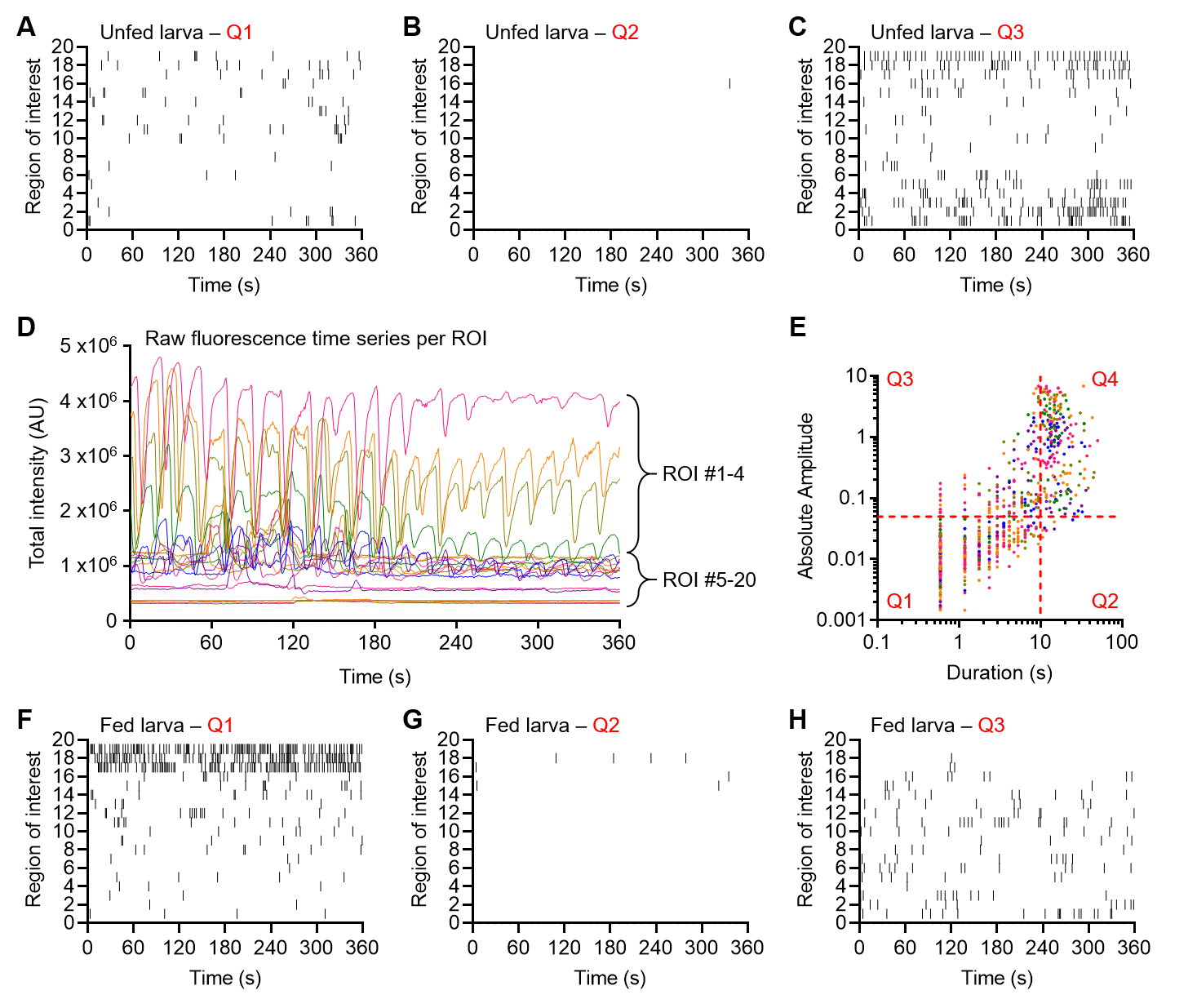

### Supplemental Table 1

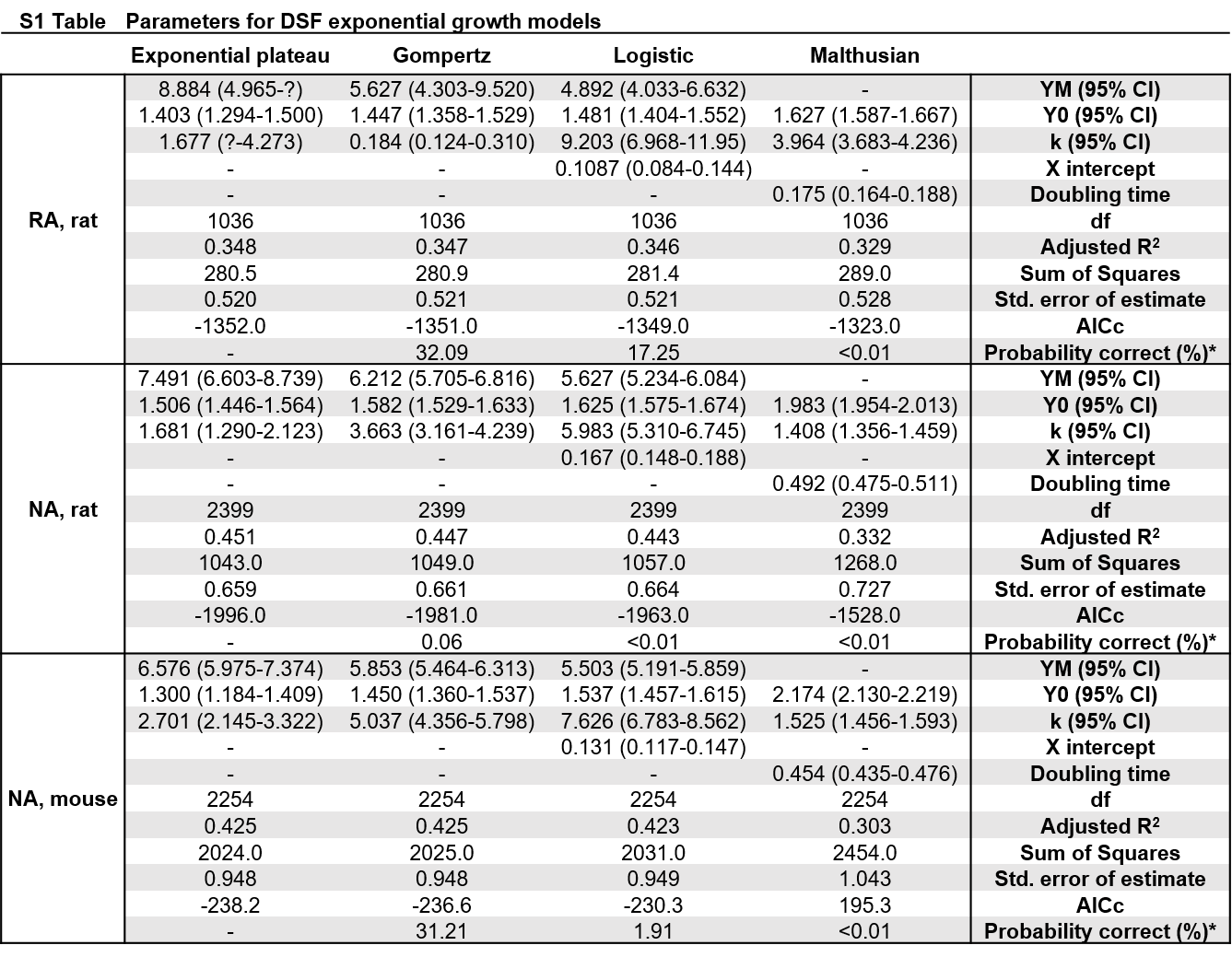
